# CRISPR/Cas9-based knockout screening revealed GSK3β as a regulator of axon initial segment structural plasticity

**DOI:** 10.64898/2026.05.21.726787

**Authors:** Yixuan Du, Ryo Egawa, Ryota Adachi, Kenta Motohara, Kazuki Furumichi, Ryota Fukaya, Hesheng Chen, Hiroshi Kuba

## Abstract

The axon initial segment (AIS) undergoes structural plasticity to tune neuronal excitability, yet the underlying molecular mechanisms remain unclear. Here we developed an in vivo CRISPR/Cas9 knockout platform using an all-in-one triple-guide RNA vector introduced via electroporation and employed this approach to identify molecules that regulate developmental AIS shortening in the chicken nucleus magnocellularis. We targeted fourteen molecules associated with microtubules and found that knockout of either glycogen synthase kinase 3β (GSK3β) or Tau impaired the AIS shortening. Conversely, overexpression of a constitutively active form of GSK3β facilitated the AIS shortening in vivo. This GSK3β-induced shortening was reproduced in slice cultures and suppressed by microtubule stabilization. Together, these findings identify GSK3β-dependent microtubule remodeling as a mechanism underlying developmental AIS shortening and establish an in vivo genetic approach for molecular screening in chick embryos.

## Introduction

The axon initial segment (AIS) is an excitable axonal domain located near the soma and serves as the site of action potential initiation^1^. This high excitability is supported by a specialized molecular organization, including the dense clustering of voltage-gated Na^+^ (Nav) channels, which are anchored via the scaffold protein, ankyrinG (AnkG), to the submembranous actin–spectrin cytoskeleton and bundled microtubules^2^. AIS structure is plastic; its length and position relative to the soma undergo activity-dependent remodeling, allowing neurons to fine-tune their intrinsic excitability and maintain stable network function^3,4^. In addition, abnormalities of AIS structure are associated with several neuropsychiatric disorders^5,6^. Therefore, elucidating the molecular mechanisms of AIS plasticity is crucial for understanding the regulation of neuronal excitability under physiological and pathological conditions.

The nucleus magnocellularis (NM), an avian homologue of the mammalian anteroventral cochlear nucleus (**Fig. 1A**), provides a well-established model for studying AIS plasticity in the developmental period. After hearing onset around embryonic day 12 (E12)^7^, NM neurons decrease the length of the AIS depending on auditory inputs, almost reaching the mature form by E21, just before hatching^8^ (**Fig. 1B**). This developmental shortening of the AIS helps tune neuronal output and preserve the fidelity of auditory signaling^9,10^. Recent work has revealed that microtubule dynamics contribute to this structural regulation of the AIS^11^, but its detailed molecular mechanisms remain elusive. An efficient and reliable loss-of-function approach for candidate molecules would be valuable, although such tools have not been available in NM neurons.

**Figure 1.**
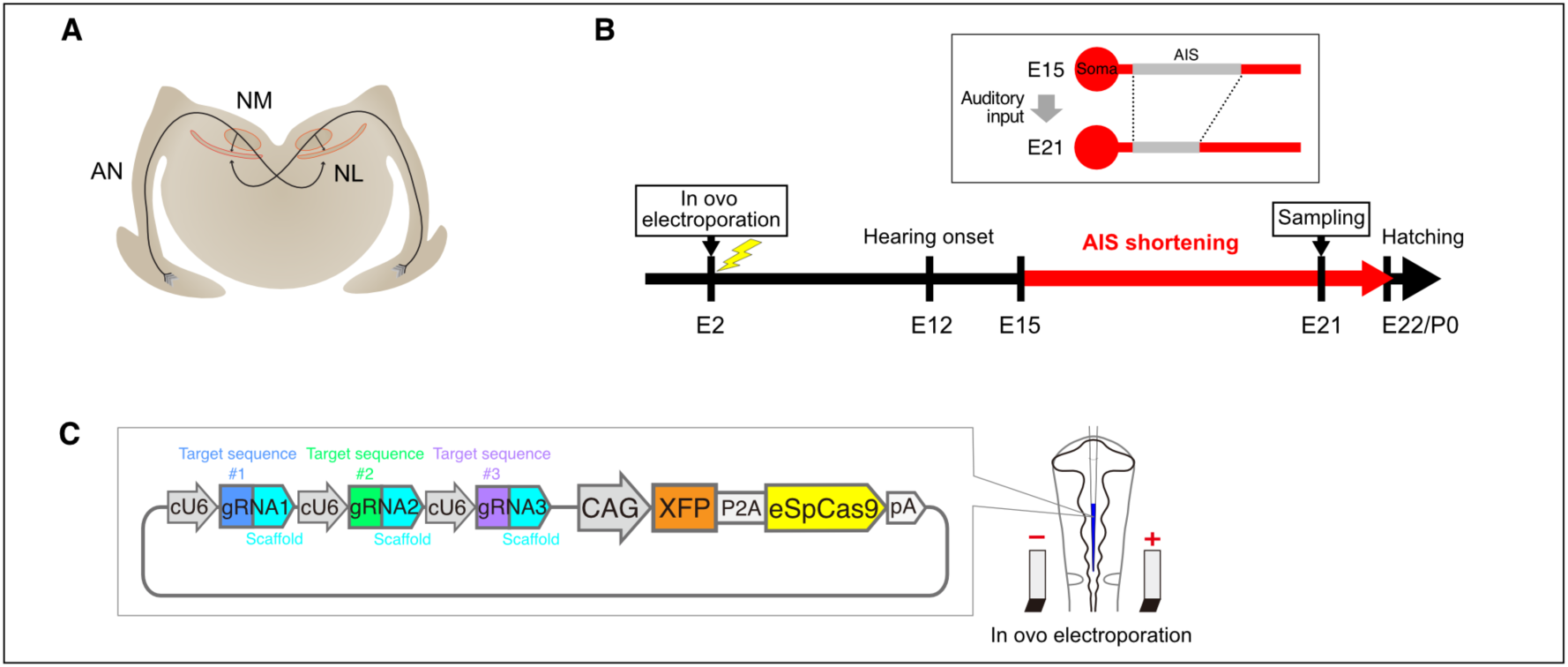
Knockout strategy for identifying regulators of AIS shortening in chick NM neurons. ***A***, Brainstem auditory circuit of the chicken. AN, auditory nerve; NM, nucleus magnocellularis; NL, nucleus laminaris. ***B***, Time course of AIS shortening in NM neurons and experimental timeline. AIS shortening begins around E15 and becomes apparent by E21. Plasmids were introduced by in ovo electroporation at E2, and brainstems were sampled at E21. ***C***, Design of the all-in-one CRISPR/Cas9 vector. Three distinct gRNAs targeting a single gene of interest are expressed under chick U6 (cU6) promoters. Simultaneously, the fluorescent reporter XFP, either tdTomato or mRuby2_smFP-Myc, and eSpCas9(1.1) are expressed under the CAG promoter. The vector was injected into the neural tube and introduced into the progenitors of NM neurons by electrical pulses (see Methods).

CRISPR/Cas9-mediated genome editing offers a promising strategy for targeted gene knockout (KO)^12,13^, and has been applied to chick embryos using in ovo electroporation^14–16^. In this approach, Cas9 nuclease is guided by a short guide RNA (gRNA) to a specific genomic locus, where it introduces a double-strand break (DSB) that is repaired mainly through non-homologous end-joining. This error-prone repair process often generates small insertions or deletions (indels) that can introduce frameshift mutations and disrupt the open reading frame of the target gene. However, DSB repair outcomes vary stochastically across individual cells, and some edited alleles may retain the reading frame through in-frame mutations^17^ or preserve partial protein function via illegitimate translation^18,19^. Such variability can reduce the reliability of in vivo knockout analyses and complicate interpretation of cellular phenotypes.

Here we established an optimized all-in-one CRISPR/Cas9 approach for in vivo gene KO in chick embryos based on a triple-target strategy^20^. Introduction of the KO vector to NM neurons via in ovo electroporation enabled loss-of-function screening for molecules involved in developmental AIS shortening. Screening of microtubule-associated molecules identified glycogen synthase kinase 3β (GSK3β), a kinase that phosphorylates microtubule-associated proteins, as a regulator of AIS shortening. Our findings thus provide a molecular mechanism linking AIS plasticity to microtubule dynamics.

## Results

### Development of an all-in-one CRISPR/Cas9 vector for KO screening

To explore the molecular mechanisms underlying AIS plasticity in the chick brainstem auditory circuit, we first designed an in vivo gene KO strategy. In ovo electroporation targeting the rhombomeres 5–8 at Hamburger–Hamilton (HH) stages 11–12 (embryonic day 2, E2) enables plasmid delivery to NM neurons^11,21^ (**Fig. 1B**).

To achieve robust disruption of target genes in the transfected neurons, we constructed an all-in-one CRISPR/Cas9 vector optimized for the chick system (**Fig. 1C**). The vector was designed to express three gRNAs targeting different sites within the same gene, based on a triple-targeting CRISPR strategy that improves KO efficiency^20^. Each gRNA was driven by the chick U6 promoter (cU6) to enhance transcriptional activity in avian cells^14^. To reduce potential off-target editing, we incorporated eSpCas9(1.1), a high-fidelity Cas9 variant^22^. In addition, a fluorescent reporter sequence encoding either tdTomato or mRuby2_smFP-Myc, a 10× Myc-tagged mRuby2^23^, was included to enable clear visualization of transfected neurons. Candidate gRNAs for each target gene were selected with reference to the predicted on-target activity scores calculated by the DeepHF algorithm^24^ (http://www.DeepHF.com/; **Table 1**). To facilitate vector construction, the three gRNA expression cassettes were integrated into a single plasmid using Golden Gate cloning^25^, allowing rapid assembly of customized KO constructs (**Supplementary Fig. 1**).

**Table 1.**
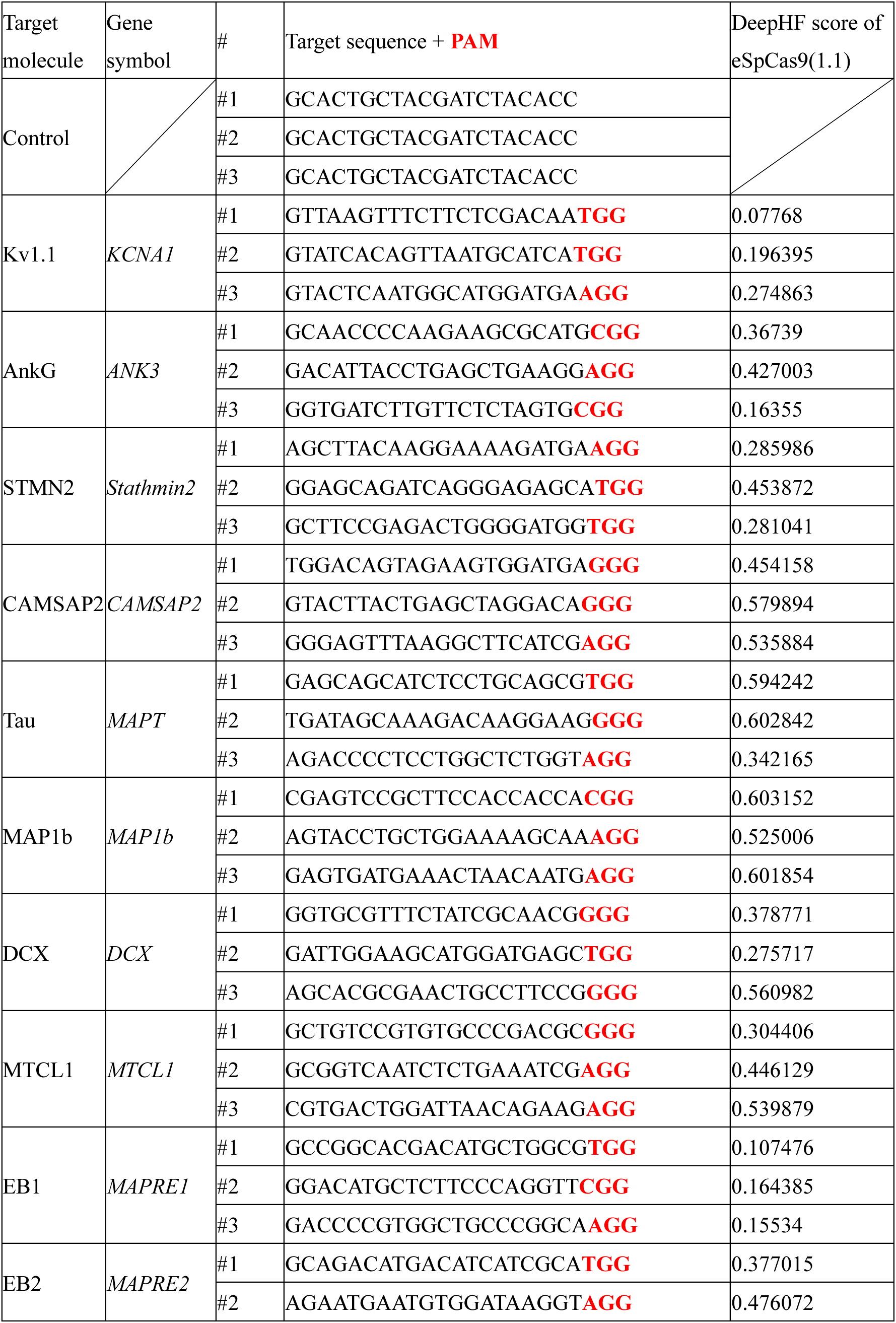
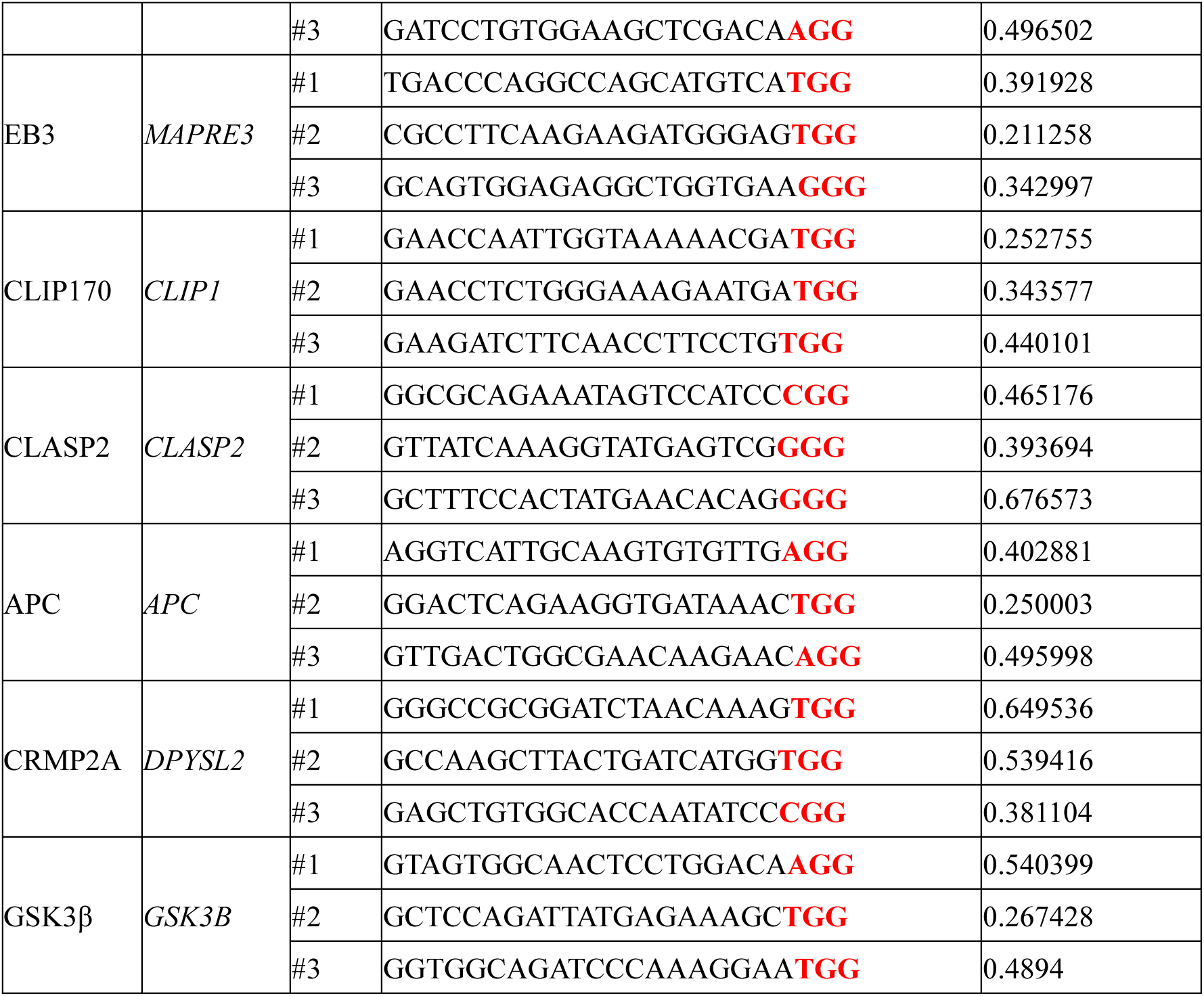
gRNA target sequences and DeepHF scores.

To validate the KO efficiency of the optimized CRISPR/Cas9 vector, we first generated a vector targeting Kv1.1, a low-voltage-activated (LVA) potassium channel abundantly expressed in NM neurons^26^. In ovo electroporation of the vector at E2 generated fluorescently labeled neurons throughout the NM region (**Fig. 2A**). These neurons showed faint or undetectable somatic Kv1.1 immunosignals compared with unlabeled neurons, or neurons transfected with a control vector expressing three non-targeting gRNAs with no predicted target sites in the chicken genome (**Fig. 2B**). The Kv1.1 signal intensity normalized to background was reduced from 4.37 ± 0.27 in control neurons to 1.56 ± 0.09 in Kv1.1-targeted neurons (p < 0.0001) (**Fig. 2C**). We next examined the electrophysiological phenotype of the Kv1.1-targeting vector using whole-cell patch-clamp recordings. Potassium currents were evoked by voltage steps from −107 mV to +13 mV at the holding potential of −77 mV in ipsilateral transfected neurons and contralateral control neurons (**Fig. 2D, E**). In control neurons, prominent tail currents appeared at the end of depolarizing pulses above ∼−70 mV (**Fig. 2D**). An LVA component that appears below –42 mV (arrow in **Fig. 2D–H**) likely reflected the Kv1 current, while a larger depolarization induced both LVA and high-voltage-activated (HVA) Kv3 currents^27^. In the transfected neurons, we also observed prominent tail currents (**Fig. 2E**). However, the Kv current at –42 mV was reduced to 34% of control levels (**Fig. 2F**), suggesting a reduction in the Kv1 current.

**Figure 2.**
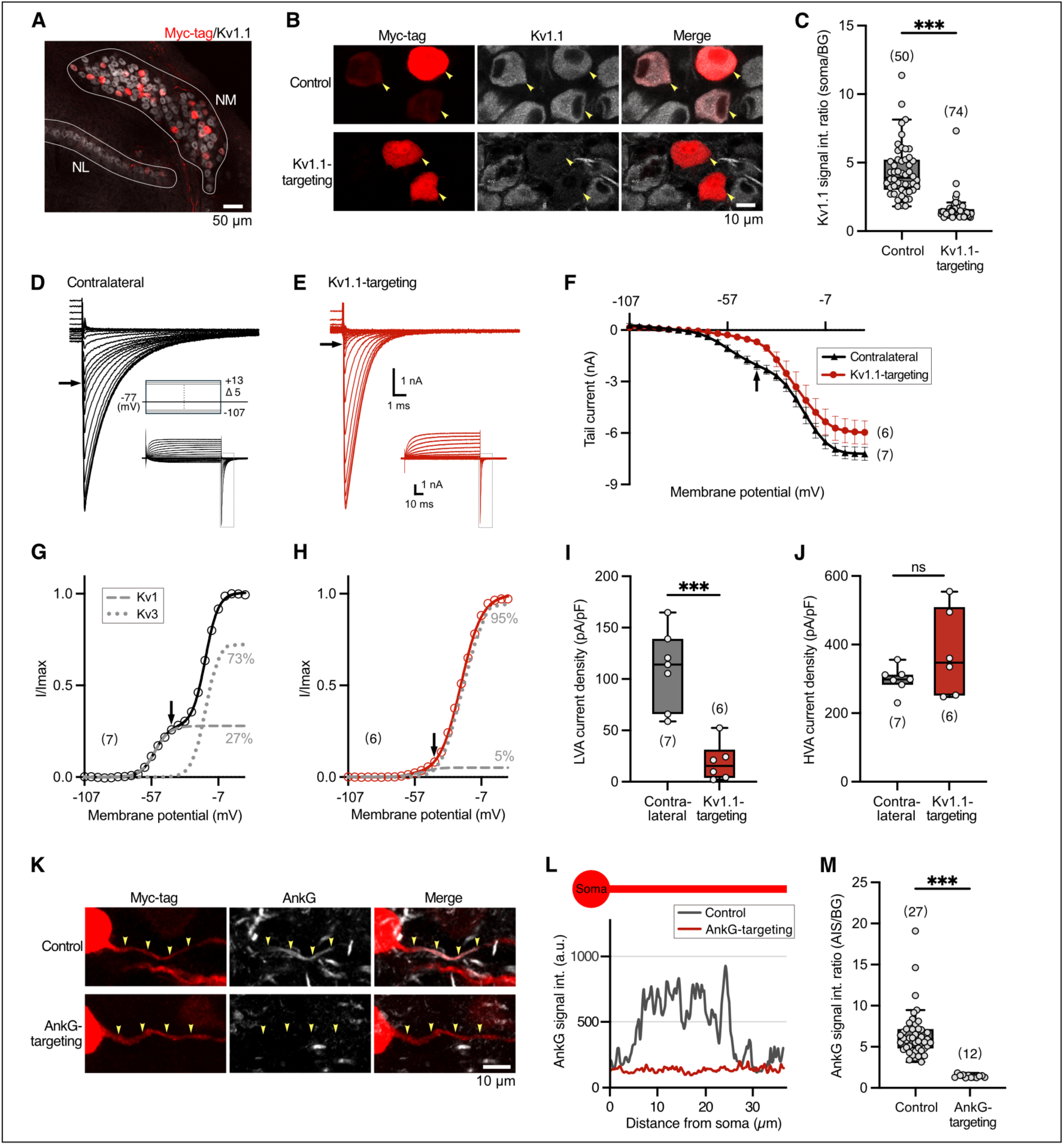
Robust knockout by all-in-one CRISPR/Cas9 vectors targeting Kv1.1 or AnkG in NM neurons. ***A***, NM neurons were stained with anti-Kv1.1 antibody (gray) and the neurons transfected with the all-in-one CRISPR/Cas9 vector were identified by the expression of Myc-tagged fluorescent reporter (red). NM and NL regions are outlined. ***B***, Kv1.1 immunoreactivity at the soma (gray) in NM neurons transfected with non-targeting control or Kv1.1-targeting vectors (red, arrowheads). ***C***, Kv1.1 somatic signal intensity. Values from individual cells are plotted as gray circles (same convention used in subsequent figures). ***D***,***E***, Whole-cell potassium tail currents recorded from contralateral control neurons (***D***, black) and ipsilateral Kv1.1-targeted neurons (***E***, red). Potassium currents were recorded using voltage steps from −107 to +13 mV in 5 mV increments from a holding potential of −77 mV. ***F***, Current-voltage relationship of tail currents. ***G***,***H***, Voltage dependence of activation curves from (***F***). The curves were fitted by a double Boltzmann equation, showing a low-voltage-activating Kv1 component (gray dashed line) and a high-voltage-activating Kv3 component (gray dotted line) (see Methods). The fraction of each component is indicated on the right. Vertical and horizontal arrows indicate −42 mV in (***D–H***). ***I***,***J***, Potassium current density (peak current amplitude normalized to membrane capacitance). (***I***) LVA current; (***J***) HVA current. Neither V_1/2_ nor slope factor differed between two groups. For the Kv1 current, V_1/2_ was −55.65 ± 2.10 mV (control) and −54.31 ± 4.09 mV (Kv1.1-targeted) (*p* = 0.78), and the slope factor was 5.16 ± 0.21 mV and 2.91 ± 1.07 mV, respectively (*p* = 0.12). For the Kv3 current, V_1/2_ was −18.45 ± 1.87 mV (control) and −21.10 ± 3.47 mV (Kv1.1-targeted) (*p* = 0.52), and the slope factor was 5.16 ± 0.14 mV and 5.45 ± 0.26 mV, respectively (*p* = 0.21). ***K***, AnkG signal at the AIS (gray, arrowheads) in NM neurons transfected with non-targeting control or AnkG-targeting vectors. ***L***, Intensity profile of AnkG signal along the axon. ***M***, AIS AnkG signal intensity. Numbers in parentheses indicate the number of cells. Biological replicates were as follows: ***C***, control, N = 3, Kv1.1-targeting, N = 3; ***F–J***, contralateral, N = 5, Kv1.1-targeting, N = 4; ***M***, control, N = 3, AnkG-targeting, N = 3. Data are mean ± SEM. \*\*\**p* < 0.001 by Welch’s t-test.

Fitting of the voltage-dependent activation curves with a double Boltzmann equation (see Methods) helped us to dissect the LVA (V₁/₂ of ∼−55 mV) and HVA (V₁/₂ of ∼−20 mV) components. The LVA component in the total current was 26.62 ± 3.08% in control neurons and 5.44 ± 2.54% in transfected neurons (**Fig. 2G, H**). A similar reduction was observed in the current density for LVA component (**Fig. 2I**), while no detectable change was observed for HVA component (**Fig. 2J**), confirming that the Kv1.1-targeting vector induces efficient KO of Kv1.1.

We then tested whether this KO system could be applied to AIS-associated proteins by targeting AnkG, a scaffold protein required for AIS organization. The AnkG immunoreactivity was detected at the AIS in all control neurons examined (27 cells), whereas it was reduced to background levels in all neurons transfected with the AnkG-targeting vector (12 cells) (**Fig. 2K–M**), indicating efficient KO of AnkG in NM neurons. Together, these results demonstrate that the optimized all-in-one CRISPR/Cas9 vector enables efficient disruption of target genes in NM neurons and is suitable for KO screening of AIS regulators in vivo.

### KO screening identified GSK3β as a potential regulator of AIS shortening

The developmental AIS shortening in NM neurons is driven by cyclin-dependent kinase 5 (CDK5)/p35-dependent microtubule destabilization^11^, suggesting that downstream microtubule-associated proteins (MAPs) may contribute to this process. We therefore used the all-in-one CRISPR/Cas9 vector to screen fourteen candidate genes associated with microtubule regulation. Six MAPs (Tau, MAP1B, DCX, CRMP2, STMN2 and CLASP2) were selected as reported CDK5 substrates^28^. To cover MAPs with distinct functions, we also included a minus-end stabilizer (CAMSAP2), a microtubule cross-linker (MTCL1), and plus-end tracking proteins (+TIPs; EB1, EB2, EB3, APC and CLIP170)^29,30^. In addition, GSK3β, a kinase that phosphorylates multiple MAPs and regulates microtubule dynamics^31^, was also included (**Fig. 3A**).

**Figure 3.**
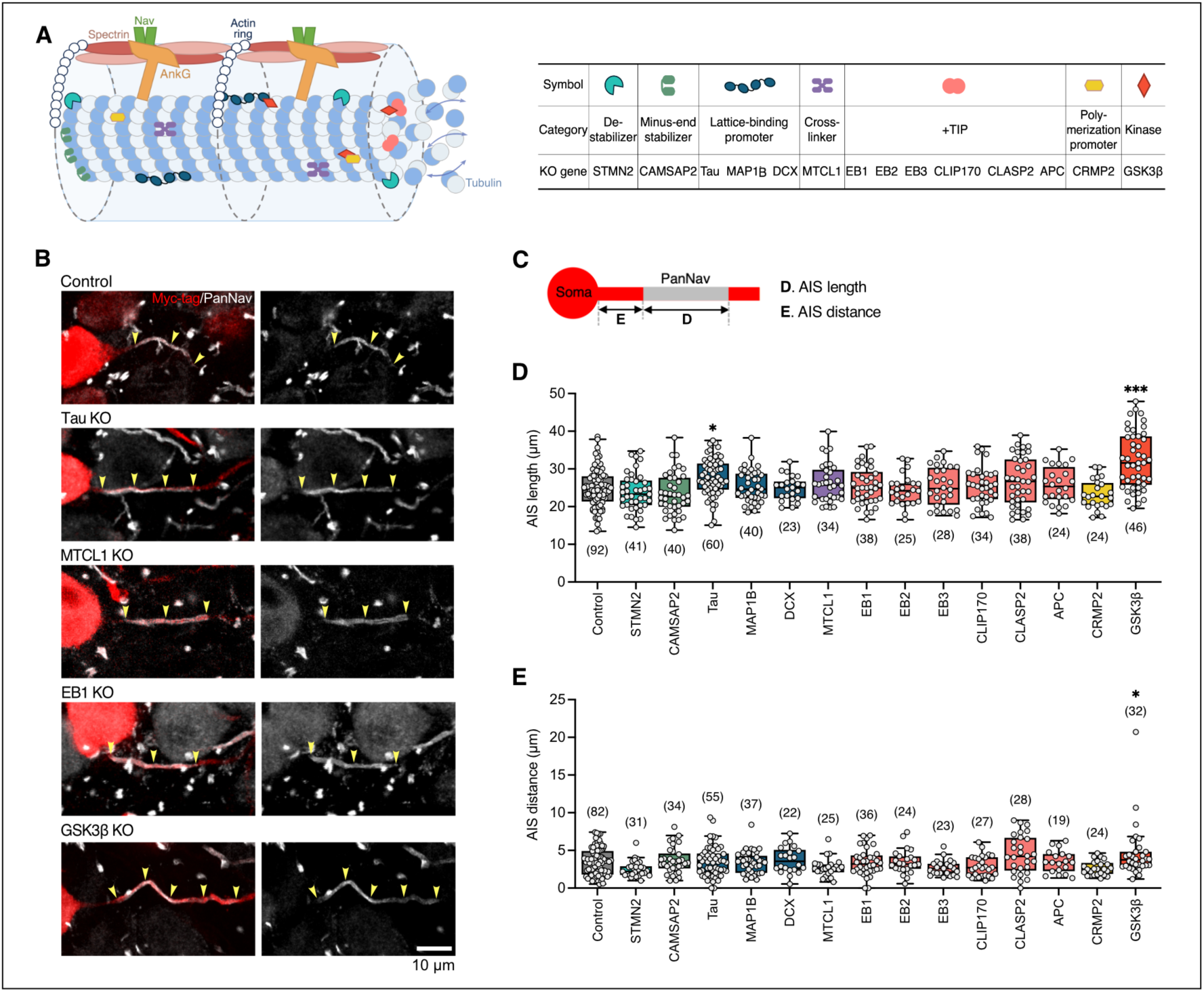
Knockout screening for microtubule regulators involved in AIS shortening. ***A***, Schematic overview of microtubule regulators at the AIS and their classification. Symbol colors for each category correspond to the colors shown in (***D***,***E***). ***B***, Representative images of AIS from NM neurons. The AIS was labeled by panNav immunostaining (gray; arrowheads), and transfected neurons were identified by Myc-tag immunostaining (red). ***C***, Definition of AIS length (***D***) and distance from the soma (***E***). ***D***, Effect of KO of candidate genes on the AIS length. ***E***, Effect of KO of candidate genes on AIS distance from the soma. Numbers in parentheses indicate the number of cells. Biological replicates were as follows: ***D***,***E***, control, N = 6; STMN2, N = 3; CAMSAP2, N = 4; Tau, N = 4; MAP1B, N = 3; DCX, N = 3; MTCL1, N = 4; EB1, N = 3; EB2, N = 3; EB3, N = 3; CLIP170, N = 3; CLASP2, N = 3; APC, N = 3; CRMP2, N = 3; GSK3β, N = 3. Data are mean ± SEM. \**p* < 0.05; \*\*\**p* < 0.001 by Dunnett’s multiple comparison test.

We assessed the effects of KOs of these candidates by measuring panNav-immunolabeled AIS signals in Myc-tag-positive neurons at E21 (**Fig. 1B, 3B**). The length and the distance of the AIS (**Fig. 3C**) were quantified (see Methods) in neurons transfected with each gene-targeting vector and compared with those in neurons expressing the non-targeting control vector. Among the candidates tested, KO of twelve of the fourteen candidates did not significantly alter either the AIS length or distance (**Fig. 3B, D, E**). These included MTCL1, EB1 and EB3, which have been implicated in AIS structural regulation in other neuronal types^32,33^. By contrast, GSK3β KO caused prominent AIS elongation (∼7 µm), accompanied by a slight distal shift (∼1 µm) of the distance from the soma (**Fig. 3D, E**). Tau KO also caused a moderate increase in the AIS length (∼3 µm) without significant change in the distance (**Fig. 3D, E**). These findings suggest that GSK3β and Tau are candidate regulators of the developmental shortening of the AIS.

### GSK3β activation facilitated AIS shortening in vivo and in vitro

To test whether increased GSK3β activity can promote AIS shortening, we introduced constructs expressing a constitutively active mutant of GSK3β (caGSK3β; S9A) together with tdTomato into NM neurons by in ovo electroporation at E2. caGSK3β expression was controlled by a Tet-On system (**Fig. 4A**) and induced by administering doxycycline (DOX, 2 μM) at E15, when the AIS shortening begins^8^ (**Fig. 4B**). At E21, caGSK3β-expressing neurons showed AISs that were ∼4 µm shorter than those of neurons expressing tdTomato alone, hereafter referred to as mock neurons (**Fig. 4C, D**), with no detectable change in the distance from the soma (**Fig. 4C, E**). These results suggest that enhanced GSK3β activity accelerated the natural course of AIS shortening during development in vivo. We also examined the effects of caGSK3β on the AIS structure in slice cultures of the chick brainstem, in which afferent input is totally disrupted^34^. Slices were prepared at E11 (0 DIV), before the onset of auditory input, from the embryos electroporated with either caGSK3β or mock constructs, and were incubated in 1× [K⁺] medium (5.3 mM) until 10 DIV (**Fig. 4A, F**). DOX treatment at 6 DIV resulted in a shorter AIS (by ∼5 μm) in caGSK3β-expressing neurons than in mock neurons (**Fig. 4G, J**). These data indicate that increased GSK3β activity promoted the AIS shortening both in vivo and in vitro. Importantly, caGSK3β-induced AIS shortening was suppressed when the cultures were treated with a microtubule-stabilizing agent, taxol (50 nM) (**Fig. 4H–J**), in agreement with the involvement of microtubule disassembly in the AIS shortening^11^.

**Figure 4.**
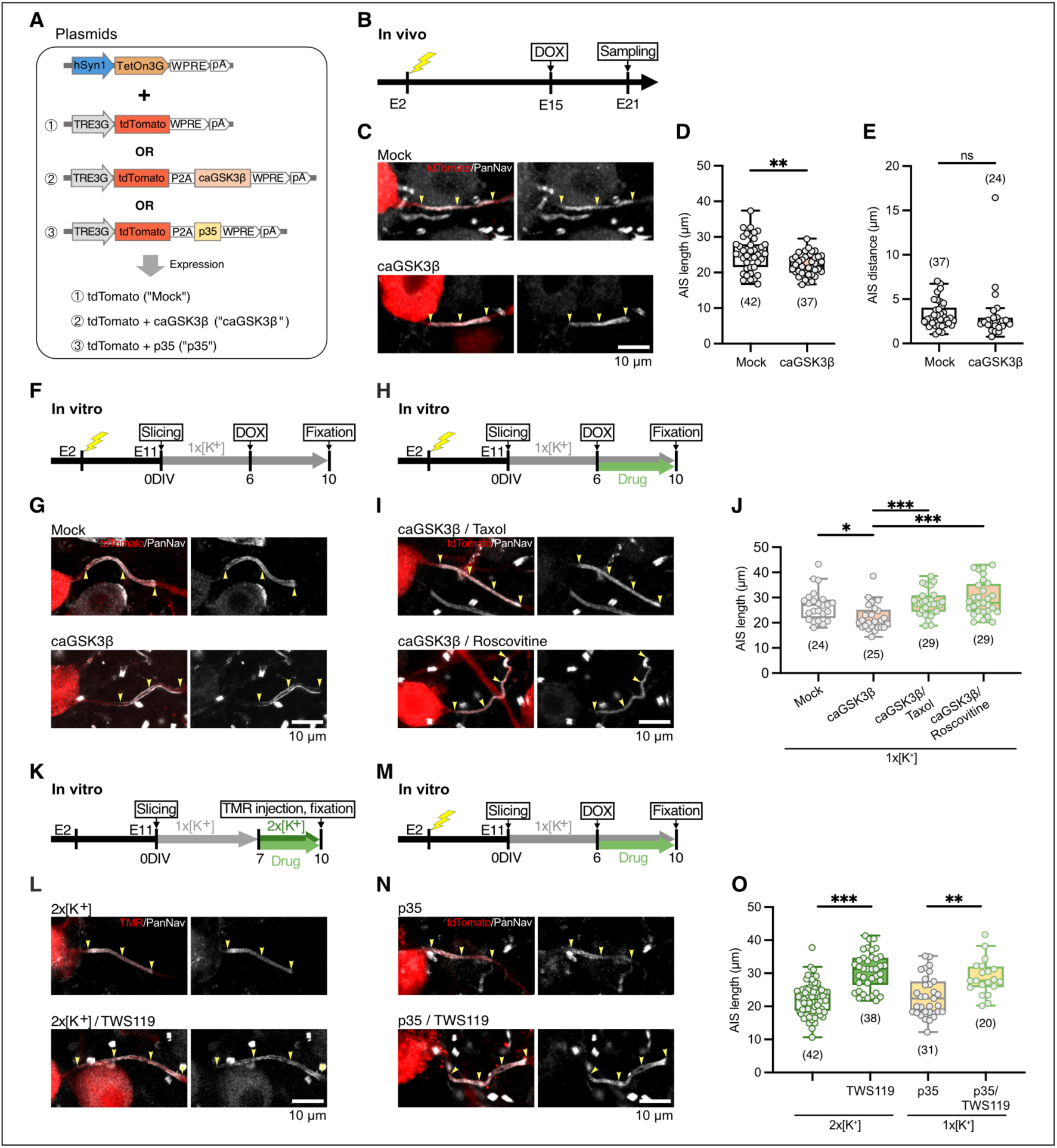
GSK3β activity facilitates AIS shortening in vivo and in vitro. ***A***, Tet-On plasmids used for inducible gene expression. ***B–E***, Increasing GSK3β activity in vivo. (***B***) Experimental timeline: mock or caGSK3β plasmids were electroporated at E2; DOX was added at E15 to induce gene expression; brainstems were sampled at E21. (***C***) AIS (gray) of NM neurons transfected with mock or caGSK3β plasmids, identified by immunostaining of tdTomato (red). (***D***,***E***) Quantification of AIS length (***D***) and distance (***E***). ***F–J***, Increasing GSK3β activity in vitro. (***F***,***H***) Experimental timeline: mock or caGSK3β plasmids were electroporated at E2; slices were prepared at E11 and cultured in 1× [K^+^] medium; DOX was added at 6 DIV to induce gene expression, either without (***F***) or with (***H***) drug treatment. (***G***) AIS of NM neurons transfected with mock or caGSK3β plasmids without drug treatment. (***I***) AIS of NM neurons transfected with caGSK3β plasmids with drug treatment. Drugs used are shown above the images. (***J***) Length of AIS. ***K–O***, Inhibiting GSK3β activity in vitro. (***K***) Experimental timeline for GSK3β inhibition under high-activity conditions shown in (***L***): slices were prepared at E11 and cultured in 1× [K^+^] medium, then switched to 2× [K^+^] medium at 7 DIV with or without TWS119 treatment. NM neurons were retrogradely labeled with dextran TMR before fixation at 10 DIV. (***L***) AIS of NM neurons cultured in 2× [K^+^] medium without (top) or with (bottom) TWS119 treatment. TMR-labeled neurons are shown in red. (***M***) Experimental timeline for GSK3β inhibition under CDK5-active conditions shown in (***N***): p35 plasmids were electroporated at E2; slices were prepared at E11 in 1× [K^+^] medium; DOX was added at 6 DIV to induce p35 expression, with or without TWS119 treatment. (***N***) AIS of NM neurons expressing p35 without (top) or with (bottom) TWS119 treatment. (***O***) Length of AIS. Arrowheads indicate AIS. The data for the 2× [K^+^] and p35 groups in (***O***) were re-plotted from previously published data in ref. 11. All other data were generated in this study. Numbers in parentheses indicate the number of cells. Biological replicates were as follows: ***D***,***E***, mock, N = 3, caGSK3β, N = 3; ***J***, mock, N = 8; caGSK3β, N = 8; caGSK3β/Taxol, N = 9; caGSK3β/Roscovitine, N = 6; ***O***, 2× [K⁺], N > 7; TWS119 in 2× [K⁺], N > 5; p35, N = 16; p35/TWS119, N = 10. Data are mean ± SEM. \**p* < 0.05; \*\**p* < 0.01; \*\*\**p* < 0.001 by Bonferroni-corrected Mann–Whitney *U*-tests.

Our previous study showed that CDK5 activation contributes to AIS shortening in the cultured NM neurons^11^. Given the known functional link between CDK5 and GSK3β in the regulation of microtubule-associated proteins^28,35^, we next examined their functional relationship in the AIS shortening using pharmacological inhibitors of these kinases in slice cultures. The AIS shortening was induced by treating slices with high K⁺ medium (10.6 mM, 2× [K⁺])^11^ (**Fig. 4K**), which was suppressed by TWS119 (0.2 µM), a GSK3β inhibitor (**Fig. 4K, L, O**). TWS119 also suppressed the AIS shortening induced by overexpression of p35^11^, an activator of CDK5 (**Fig. 4M–O**). These results indicate that GSK3β activity is required for the AIS shortening induced by CDK5 activation. Importantly, the AIS shortening induced by caGSK3β was suppressed by a CDK5 inhibitor, roscovitine (2 µM) (**Fig. 4H–J**), suggesting that CDK5 activity is also required under GSK3β activation for AIS shortening. Together, these results suggest that GSK3β and CDK5 act in a shared microtubule-dependent mechanism to regulate AIS shortening, rather than through a pathway mediated by GSK3β alone.

## Discussion

In this study, we established an optimized all-in-one CRISPR/Cas9 KO system for chick embryos, achieving efficient gene disruption by using in ovo electroporation. This approach allowed us to screen for candidate regulators of AIS shortening that had been implicated in microtubule remodeling. Unexpectedly, KO of most classical MAPs, except Tau, exerted no detectable effect on the AIS shortening. In contrast, KO of GSK3β suppressed the AIS shortening, whereas its activation promoted this process. These findings suggest that GSK3β acts as a positive regulator of activity-dependent AIS shortening in NM neurons.

### An all-in-one triple-target CRISPR/Cas9 system for gene disruption in chick embryos

A methodological feature of this study is the development of a CRISPR/Cas9-based gene editing system for chick embryos that improves the correspondence between fluorescent labeling and gene disruption at the single-cell level. In previous in vivo applications of CRISPR/Cas9, particularly those using separate vectors for genome editing and visualization, reporter expression did not necessarily indicate successful disruption of the target gene in the labeled cells, because the two kinds of vectors could be delivered in a mosaic manner across cells^36–38^, and the stochastic nature of DSB repair could result in partial preservation of gene function in individual neurons^17–19^. These issues are particularly relevant when analyzing cellular and subcellular phenotypes, such as AIS morphology, where accurate identification of knockout cells is essential. To address these issue, we developed an all-in-one vector that ensured co-expression of Cas9 nuclease, three gRNAs, and a fluorescent reporter in the same cells. This design increases the correspondence between fluorescent labeling and delivery of the gene-editing machinery. The use of three gRNAs was intended to reduce the likelihood of preserved target gene function, while eSpCas9(1.1) and score-guided gRNA selection were incorporated to limit potential off-target editing (**Fig. 1C**). Consistent with this design, Kv1.1 targeting reduced both Kv1.1 immunoreactivity and low-voltage-activated potassium currents in reporter-positive NM neurons (**Fig. 2B–J**), whereas AnkG targeting abolished AIS AnkG immunoreactivity (**Fig. 2K–M**). These validation experiments support the use of this system for single-cell loss-of-function analysis in chick NM neurons.

This system should also be applicable to other studies that require analysis of individually labeled cells after gene disruption in embryonic chickens. Chick embryos are widely used for in vivo studies of developmental patterning, organogenesis, enhancer activity, and preclinical oncology using the chick chorioallantoic membrane^39,40^, many of which rely on local in ovo electroporation for spatially restricted gene manipulation^41–43^. In such studies, a fluorescent reporter is often used to identify manipulated cells, but it does not necessarily indicate effective gene disruption in the same cell. By linking fluorescent labeling with delivery of the gene-editing components, our all-in-one system may therefore facilitate reliable single-cell or subcellular phenotyping after gene disruption. In addition, the streamlined Golden Gate-based cloning strategy (**Supplementary Fig. 1**) allows the system to be readily applied to different target genes and experimental paradigms, thereby providing a versatile platform for molecular screening in chick embryos.

A potential limitation of single-gene KO approaches is that they may trigger genetic compensation^44^. Mutations introduced by Cas9 can induce nonsense-mediated decay of mutant mRNAs, which may in turn upregulate the expression of functionally related paralogs as a compensatory response^45^. Such compensation could mask phenotypes, especially for genes with overlapping functions. Multiplex KO of paralogous genes may help address this issue. However, simply adding more promoter-driven gRNA expression cassettes may not be optimal, as transcriptional interference could reduce the expression of individual cassettes^46–48^. Further development of this system could therefore incorporate alternative gRNA expression strategies, such as polycistronic tRNA–gRNA systems, which enable the production of multiple gRNAs from a single transcript with less transcriptional interference^49–51^.

### Roles of GSK3β and CDK5 in AIS shortening

We previously showed that activation of CDK5 promotes microtubule disassembly at the distal AIS and facilitates the AIS shortening in NM neurons^11^. In this study, we identified GSK3β as a regulator that functionally cooperates with CDK5 in this type of plasticity (**Fig. 4H–J, M–O**), reminiscent of the contribution of GSK3β to AIS structure in dissociated hippocampal neurons^52^. How, then, could CDK5 and GSK3β be recruited for the AIS shortening?

Structural AIS plasticity is triggered by Ca²⁺ influx^4,53^, and CDK5 activation downstream of Ca²⁺ influx contributes to AIS shortening in NM neurons^11^. Ca^2+^ influx also activates calcineurin (PP2B), which can dephosphorylate GSK3β at Ser9 and thereby enhance its kinase activity^54^. Indeed, calcineurin has been implicated in AIS remodeling in dissociated hippocampal neurons^53^ and in AIS shortening in NM neurons^11^. In parallel, Ca²⁺-dependent activation of the Ras/Raf–MEK–ERK pathway^55^ may also contribute, as this pathway can activate GSK3β by phosphorylating Tyr216^56^. Notably, ERK activation can increase CDK5 activity by promoting p35 expression via Egr1^57^, consistent with suppressed AIS shortening in NM neurons by MEK inhibition^11^. Moreover, under the condition of Ca²⁺ overload, calpain-mediated cleavage of p35 generates p25, which not only enhances CDK5 activity but also preferentially binds to and activates GSK3β^58^. These pathways provide potential routes by which Ca²⁺ signals could engage both CDK5 and GSK3β during AIS shortening. Although previous work proposed an indirect inhibitory crosstalk in which CDK5 suppresses GSK3β through PP1-dependent regulation^59^, our slice culture data, in which TWS119 suppressed p35-induced AIS shortening argue against this pathway (**Fig. 4M–O**). Instead, our results support a cooperative model in which CDK5 and GSK3β converge on shared microtubule-associated substrates, potentially through priming phosphorylation by CDK5 followed by GSK3β-dependent phosphorylation^60^. Together with these reported mechanisms, our data suggest that CDK5 and GSK3β may convert Ca^2+^-dependent signals into microtubule reorganization, thereby tuning the location of AIS and consequent excitability.

### Roles of Tau in AIS shortening

Inhibition of either GSK3β or CDK5 activity was sufficient to occlude the AIS shortening by the other, suggesting that the two kinases may share common substrates for microtubule reorganization at the AIS. Among the MAPs examined in our screening, only Tau KO affected the AIS shortening (**Fig. 3B, D**). Importantly, multiple evidence supports a functional link between Tau and the AIS. In mouse CA1 pyramidal neurons, hyperphosphorylated Tau reduces membrane excitability via AIS remodeling^61^, whereas in human iPSC-derived neurons, pathogenic Tau mutations impair activity-dependent AIS plasticity and disrupt homeostasis of network activity^62^. Moreover, Tau is regulated via hierarchical phosphorylation by both CDK5 and GSK3β, in which CDK5-mediated priming phosphorylation at Ser235/Ser404 facilitates subsequent GSK3β-dependent phosphorylation at Thr231/Ser400/Ser396, reducing the microtubule-binding affinity of Tau and promoting microtubule destabilization^60,63^. These findings suggest that Tau may be a substrate regulated by GSK3β and CDK5, enabling the Ca²⁺-dependent microtubule reorganization and AIS shortening in NM neurons. A recent study further showed that hyperactivity of GSK3β reduces Tau binding to microtubules and destabilizes microtubules at their plus ends^64^. This is consistent with the AIS shortening that proceeds at the distal end in NM neurons^8^, although the spatiotemporal regulation of GSK3β activity during activity-dependent AIS shortening remains unclear. Nevertheless, the AIS elongation after Tau KO should not be interpreted as direct evidence that Tau alone drives this process, because chronic genetic deletion may induce compensatory changes in related MAPs^65–67^.

The lack of detectable phenotype in AIS position after KO of most other MAPs may also reflect extensive functional redundancy among MAPs. Indeed, substantial functional overlap exists within the EB family and the broader +TIP network (EBs–CLIP170–CLASP2–APC)^68–71^, and strong compensatory relationships are observed between Tau and MAP1B^67,72^, and between CAMSAP2 and CAMSAP3^73^. Alternatively, the effects of MAPs on microtubule reorganization at the AIS may vary across experimental systems, species and neuronal types. Many relevant mechanisms were established in dissociated mammalian neuronal cultures^32,74–77^, whereas the avian NM neurons undergo AIS remodeling in vivo and may rely on different combinations of redundant regulators. Supporting this possibility, TRIM46 knockdown disrupts AIS microtubule organization in cultured neurons, whereas AIS formation remains largely intact in TRIM46 KO mice^75,78^.

In summary, we established an optimized all-in-one CRISPR/Cas9 KO system for single-cell loss-of-function analyses in chick embryos and identified GSK3β as a regulator of AIS shortening in NM neurons during development. Thus, our genetic strategy would help advance a comprehensive understanding of the molecular network controlling AIS plasticity.

## Methods

### Ethics statement

All animal experiments were approved by the Animal Experiment Committee of Nagoya University (approval number: M240042-001) and performed in accordance with the regulations on animal experiments at Nagoya University. The experimental protocols were carried out in accordance with the Fundamental Guidelines for Proper Conduct of Animal Experiment and Related Activities in Academic Research Institutions (Notice No. 71 of the Ministry of Education, Culture, Sports, Science and Technology, 2006), the Standards relating to the Care and Keeping and Reducing Pain of Laboratory Animals (Notice of the Ministry of the Environment No. 88 of 2006) and the Standards relating to the Methods of Destruction of Animals (Notice No. 40 of the Prime Minister’s Office, 1995).

### Animals

Chick embryos (*Gallus domesticus*) of either sex from E2 to E21 were used. Fertilized eggs were incubated in a humidified incubator at 37.5°C until reaching the desired developmental stage. Embryos were anesthetized by cooling the eggs in ice-cold water. Unless otherwise noted, brainstem tissue containing the rostral half of the NM region was used for experiments^8^.

### Plasmid construction

The all-in-one CRISPR/Cas9 vector optimized for the chick system was designed based on a multiple gRNA expression system described previously^25^. Briefly, the human U6 promoter, *Bpi*I digestion sites, and gRNA scaffolds contained in the gRNA expression cassettes of pX330A-1x3 (BB) (Addgene #58767), pX330S-2 (BB) (Addgene #58778), and pX330S-3 (BB) (Addgene #58779) were replaced with chicken U6.3 promoter^79^, *Sap*I digestion sites, and the F+E gRNA scaffold^14,80^ by In-Fusion cloning (Takara Bio). In addition, the CBh promoter in pX330A-1x3 (BB) was substituted with the CAG promoter, and the original Cas9 nuclease was replaced with a construct consisting of tdTomato or mRuby2_smFP-Myc (pCAG-mRuby2smFP-Myc; Addgene #71816)^23^, a P2A self-cleaving peptide sequence^81^, and eSpCas9(1.1) (Addgene #71814)^22^. To facilitate the expression check after electroporation, the inactivated tripeptide (GGG) in the chromophore of mRuby2_smFP-Myc was reactivated by replacing it with the original tripeptide (MYG). The modified backbone plasmids of pX330A-1x3 (BB), pX330S-2 (BB), and pX330S-3 (BB), hereafter referred to as GE1-1x3 (BB), GE2 (BB), and GE3 (BB), respectively, served as templates for insertion of arbitrary target sequences. Single-plasmid vectors expressing three gRNAs per target gene were generated according to a procedure described previously^25^ (see **Supplementary Fig. 1**). In brief, forward and reverse oligonucleotides (IDT) harboring each 20-nt target sequence (**Table 1**) were annealed and ligated into *Sap*I-digested GE1-1x3 (BB), GE2 (BB), and GE3 (BB), respectively. The three resulting gRNA expression cassettes of each plasmid were then assembled into a single vector by Golden Gate cloning using *Bsa*I, thereby producing a construct carrying three tandem gRNA cassettes.

Candidate gRNA sequences for each gene were listed using CHOPCHOP v3^82^ (https://chopchop.cbu.uib.no/), restricting the search to sites beginning with GN or NG at the 5′ end. Among them, three target sequences were selected based on the following criteria: targeting functionally or structurally important domains, avoidance of low DeepHF scores, exclusion of sites with ≤2-bp off-target mismatches, and exclusion of positions overlapping known SNPs.

pTRE-tdTomato-WPRE and phSyn1-TetOn3G were described previously^11^. The pTRE-tdTomato-P2A-caGSK3β-WPRE construct was generated by inserting caGSK3β (Tag5Amyc-GSK3b CA; Addgene #16261)^83^ into pTRE3G-tdTomato-P2A-CDK5-WPRE^11^ by In-Fusion cloning (Takara Bio). All constructs were verified by Sanger sequencing.

### In ovo electroporation and incubation

The protocol was based on the previous report^11^. Briefly, a plasmid cocktail (1 μg μL⁻¹; 0.3–0.4 µl) was injected into the neural tube of the chick embryos at E2 (HH Stage 11–12)^84^ and introduced into the right side of the hindbrain (rhombomeres 5–8). Electrical pulses were delivered using an electroporator (NEPA21, NEPAGENE) with a bipolar electrode (CUY613P1, NEPAGENE). Pulse parameters were as follows. For CRISPR/Cas9 vectors: poring pulse: 15 V, 30 ms width, 50 ms interval, 3 pulses, 10% decay, and transfer pulse: 5 V, 50 ms width, 50 ms interval, 10 pulses, 20% decay; for other vectors: poring pulse, 15 V, 30 ms width, 50 ms interval, 3 pulses, 10% decay; transfer pulse, 5 V, 50 ms width, 50 ms interval, 5 pulses, 40% decay.

### Immunohistochemistry

Brainstems were dissected from embryos at E21 and fixed with periodate-lysine-paraformaldehyde (PLP) fixative (1% paraformaldehyde, 2.7% lysine HCl, 0.21% NaIO4, and 2.85 mM Na_2_HPO_4_) for 1.5 h at 4°C. For coronal sections, brainstems were cryoprotected by overnight immersion in 30% (w/w) sucrose in PBS, embedded in OCT compound, and sectioned at 50 µm thickness. Sections were incubated overnight with primary antibodies: rabbit polyclonal AnkG antibody (5 µg mL⁻¹)^85^, rabbit polyclonal Kv1.1 antibody (1.5 µg mL⁻¹, Alomone Labs), mouse monoclonal panNav antibody (5 µg mL⁻¹, Sigma-Aldrich), rat monoclonal Myc-tag antibody (2.5 µg mL⁻¹, Proteintech), and rabbit polyclonal RFP antibody (1.06 µg mL⁻¹, Rockland). Sections were then incubated with Alexa dye–conjugated secondary antibodies (10 µg mL⁻¹; Thermo Fisher Scientific) for 2 h. Antibodies were diluted in PBS containing 1% donkey serum, 0.05% carrageenan, and 0.3% Triton X-100 to reduce non-specific binding of the antibodies. After washing, sections were mounted and imaged using a confocal laser-scanning microscope (AX R, Nikon) equipped with a 60×, 0.95-NA objective (Nikon).

Serial sections were Z-stacked at a step of 0.3 µm. For AnkG and panNav measurement, 15–25 confocal planes were Z-stacked, and AIS distance from the soma and AIS length were measured as described previously^8^. For Kv1.1 measurements, images were captured from a single confocal plane, and signals were quantified within the somatic ROIs (>10 µm²) in >15 neurons per animal. For intensity quantification, images were acquired using identical microscope settings, and signal intensity was normalized to the background signal. Background signal was estimated as the average of the mean intensities of 3–5 small ROIs placed within the NM region but devoid of obvious AnkG- or Kv1.1-positive structures. Image analysis was performed using Fiji (ImageJ), and figures were assembled using Affinity (https://www.affinity.studio/).

### Electrophysiology

Slice preparation and whole-cell voltage-clamp recordings were performed following the protocol reported previously^86^. The isolated brainstem was immersed in high-glucose artificial cerebrospinal fluid (HG-ACSF; in mM: 75 NaCl, 2.5 KCl, 26 NaHCO₃, 1.25 NaH₂PO₄, 1 CaCl₂, 3 MgCl₂, and 100 glucose) continuously equilibrated with 95% O₂/5% CO₂. Tissues were embedded in 3.5% (w/v) agarose (Nacalai), and three to four coronal sections (230–250 μm thick) were cut using a vibratome (VT1200, Leica). Slices were incubated in the HG-ACSF at room temperature for ∼30 min before use. Whole-cell voltage-clamp recordings were obtained at 20°C using patch pipettes (3–4 MΩ) filled with a Cs⁺-based internal solution (in mM: 155 CsMeSO₃, 5 NaCl, 3 MgCl₂, 0.2 EGTA, and 10 HEPES adjusted with CsOH; pH 7.2). The use of Cs⁺ internal solution together with the lower recording temperature effectively reduced and slowed K⁺ currents, thereby minimizing series resistance errors and improving measurement accuracy^27,86^. Slices were continuously perfused with a normal ACSF containing the following (in mM): 125 NaCl, 7.5 KCl, 26 NaHCO_3_, 1.25 NaH_2_PO_4_, 0.5 CaCl_2_, 1 MgCl_2_, 0.2 CdCl_2_, 0.5 NiCl_2_, 17 glucose, 0.001 TTX (Tocris Bioscience), 0.02 DNQX (Tocris Bioscience), 0.01 SR95531 (Tocris Bioscience), bubbled with 95% O₂/5% CO₂. Liquid junction potentials were corrected (7.0 mV), and series resistance was compensated electronically up to 70%.

Voltage-dependent activation of the potassium current was analyzed by plotting the amplitude of the tail current after normalization and fitting the curve with a double Boltzmann equation^27,86^ as follows: I/I_max_ = A1/{1 + exp[− (V_m_ − V_1/2_1)/S1]} + A2/{1 + exp[− (V_m_ − V_1/2_2)/S2]}, where I is the tail current amplitude, I_max_ is the maximum tail current amplitude, V_m_ is the membrane potential, A1 and A2 are weighting factors, V_1/2_1 and V_1/2_2 are half-activation voltages, and S1 and S2 are slope factors of each component. The potassium current density was calculated by normalizing the current to the membrane capacitance.

### Organotypic slice culture

The detailed procedure has been previously described^11^. Briefly, brainstems were dissected as described above. Four to five coronal slices (200 µm) were prepared using a vibratome. Slices containing rostral one-third of NM regions were collected, transferred onto a Millicell membrane insert (Millipore) in a culture dish (35 mm), and cultured for 10 d in vitro (DIV) in Neurobasal medium (Thermo Fisher Scientific) containing 2% B-27 serum-free supplement (Thermo Fisher Scientific), 1 mM glutamate solution (Fujifilm), and 1% penicillin-streptomycin solution (Thermo Fisher Scientific).

During the first 4 d, 5% fetal bovine serum (Biowest) was added, and half of the medium was changed twice a week. NM neurons were depolarized for 3 d from 7DIV by adding KCl to the medium. For overexpression experiments, DOX was added to the culture medium at 6DIV. The kinase inhibitors, roscovitine (Calbiochem) or TWS119 (MedChemExpress) were added to the culture medium simultaneously with DOX. NM neurons in the culture were labeled by tetramethylrhodamine (TMR)-conjugated dextran (MW 3000; Thermo Fisher Scientific; 10–40% in 0.1 M phosphate buffer adjusted to pH 2.0 with HCl), unless tdTomato was overexpressed. Slices were fixed with PLP fixative containing 0.4% paraformaldehyde for 12 min at room temperature. Slices were incubated overnight with primary antibodies, followed by Alexa dye–conjugated secondary antibodies for 2 h at room temperature. Primary antibodies used were mouse monoclonal panNav antibody, rabbit polyclonal TRITC (TMR) (2.5 µg mL⁻¹, Thermo Fisher Scientific) antibody, and rabbit polyclonal RFP antibody. Slices were mounted and imaged using a confocal laser-scanning microscope (FV1000, Olympus) equipped with a 60×, 1.35-NA objective (Olympus). Z-stacks were acquired with a step size of 0.8 µm. Image analysis was performed as described above.

### Statistics and Reproducibility

For all quantitative analyses, each data point represents an individual neuron. The number of cells analyzed is indicated in parentheses in each graph, and the number of embryos or independent slice cultures is provided in the corresponding figure legends. Biological replicates were defined as independent embryos for in vivo experiments and as independent slice cultures prepared from different embryos for slice culture experiments. Data are presented as mean ± standard error of the mean (SEM), unless otherwise stated.

Statistical analyses were performed using SPSS (IBM). For comparisons between two groups, Welch’s *t*-test was applied without assuming equal variances. The robustness of the results was confirmed, as similar outcomes were obtained using Student’s *t*-test and the Mann–Whitney *U*-test. For comparisons between the control group and each genetically manipulated group in Fig. 3, Dunnett’s multiple comparison test was used. For comparisons among the four groups in Fig. 4J and 4O, Bonferroni-corrected Mann–Whitney *U*-tests were performed. Statistical significance was set at p < 0.05.

## Data availability

The data that support the findings of this study are available from the corresponding author upon reasonable request.

## Acknowledgments

We acknowledge the Division for Medical Research Engineering, Nagoya University Graduate School of Medicine, for access to the AX R confocal microscope and NanoDrop 2000. We also thank Feng Zhang, Takashi Yamamoto, Loren Looger and Mien-Chie Hung for providing eSpCas9(1.1) (Addgene#71814), pX330A-1x3 (BB) (Addgene#58767), pX330S-2 (BB) (Addgene#58778), pX330S-3 (BB) (Addgene#58779), pCAGmRuby2smFPMyc (Addgene#71816) and Tag5Amyc-GSK3b CA (Addgene#16261). This work was supported by grants-in-aid from MEXT (21H02577, 24H00584 to H.K.; 23K05986, 20K15915 to R.E.; 24K09658 to R.A.), JST SPRING (JPMJSP2125 to Y.D.), and Nagoya University CIBoG WISE program from MEXT to Y.D.

## Author contributions

Conceptualization, Y.D., R.E. and H.K.; Methodology, R.E., Y.D., K.M. and K.F.; Investigation, Y.D., R.E., and R.A.; Formal analysis and Data curation, Y.D., R.E. and H.C.; Visualization, Y.D. and R.E.; Writing – original draft, Y.D. and R.E.; Writing – review & editing, Y.D., R.E., R.A., R.F. and H.K.; Funding acquisition, H.K., R.E., R.A. and Y.D.; Supervision, H.K.

