## Supplementary Figure 1 for "CRISPR/Cas9-based knockout screening revealed GSK3β as a regulator of axon initial segment structural plasticity"

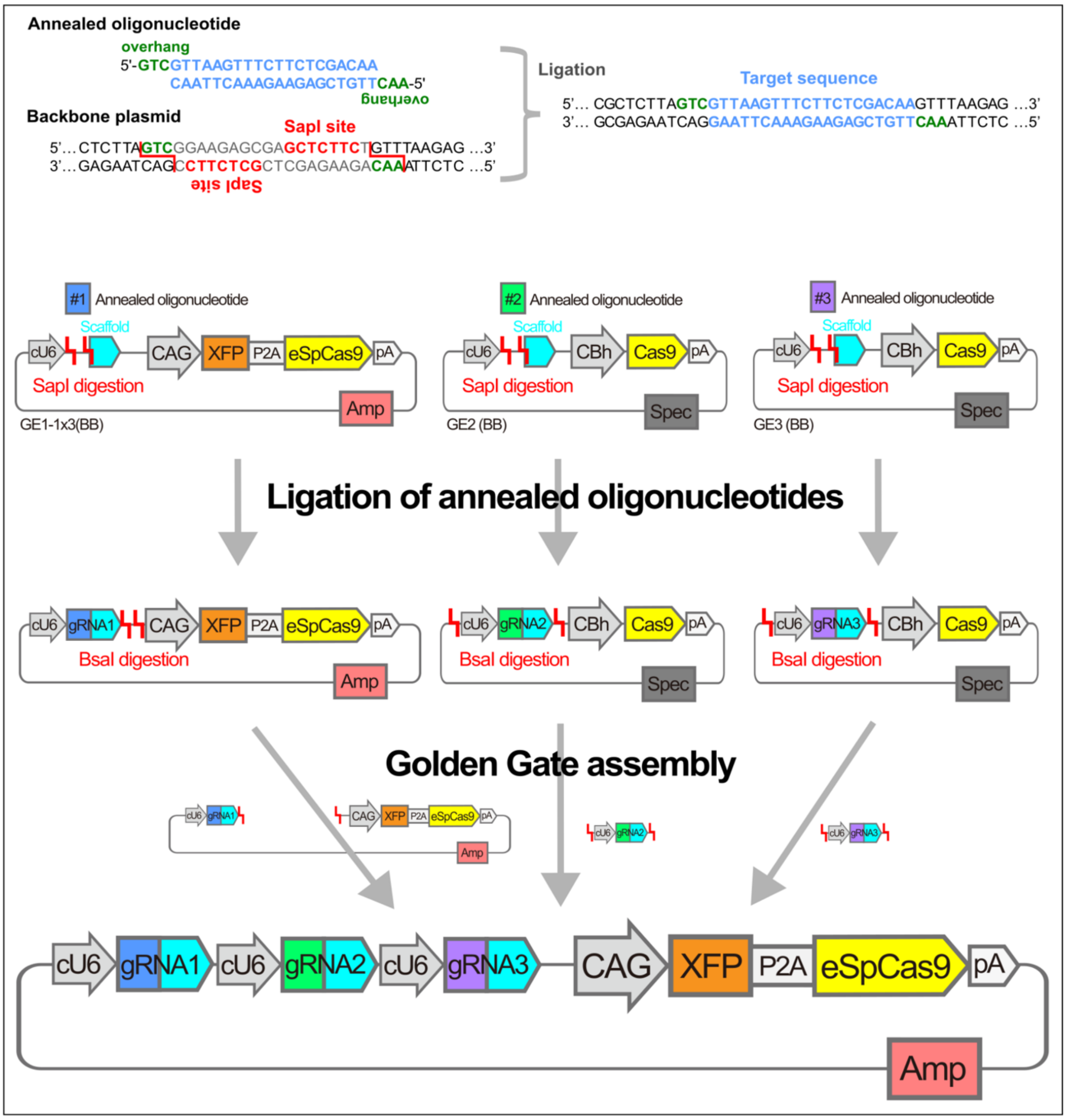


**Supplementary Figure 1. Construction of an all-in-one CRISPR/Cas9 vector expressing three distinct gRNAs.**

Based on the list of gRNA target sequences (**Table 1**), pairs of 23-nt forward and reverse oligonucleotides were designed. Each forward oligonucleotide contained a GTC overhang appended upstream of the 20-nt target sequence adjacent to the protospacer adjacent motif (PAM), whereas each reverse oligonucleotide contained an AAC overhang appended upstream of the complementary target sequence. These oligonucleotide pairs were annealed and ligated into SapI-digested backbone plasmids GE1-1x3 (BB), GE2 (BB), and GE3 (BB), respectively. The three resulting plasmids harboring individual target sequences were then combined using Golden Gate cloning, in which BsaI digestion and ligation occur simultaneously, to assemble a single vector containing three gRNA expression cassettes arranged in tandem. This procedure represents a partial modification of the multiple gRNA expression system originally described previously[25].
